# Dynamic regulation of immunity through post-translational control of defense transcript splicing

**DOI:** 10.1101/736249

**Authors:** Keini Dressano, Philipp R Weckwerth, Elly Poretsky, Yohei Takahashi, Carleen Villarreal, Zhouxin Shen, Julian I. Schroeder, Steven P. Briggs, Alisa Huffaker

## Abstract

Survival of all living organisms requires the ability to detect attack and swiftly counter with protective immune responses. Despite considerable mechanistic advances, interconnectivity of signaling circuits often remains unclear. A newly-characterized protein, IMMUNOREGULATORY RNA-BINDING PROTEIN (IRR), negatively regulates immune responses in both maize and Arabidopsis, with disrupted function resulting in enhanced disease resistance. IRR physically interacts with, and promotes canonical splicing of, transcripts encoding defense signaling proteins, including the key negative regulator of pattern recognition receptor signaling complexes, CALCIUM-DEPENDENT PROTEIN KINASE 28 (CPK28). Upon immune activation by Plant Elicitor Peptides (Peps), IRR is dephosphorylated, disrupting interaction with *CPK28* transcripts and resulting in accumulation of an alternative splice variant encoding a truncated CPK28 protein with impaired kinase activity and diminished function as a negative regulator. We demonstrate a novel circuit linking Pep-induced post-translational modification of IRR with post-transcriptionally-mediated attenuation of CPK28 function to dynamically amplify Pep signaling and immune output.

**One Sentence Summary:** Plant innate immunity is promoted by post-translational modification of a novel RNA-binding protein that regulates alternative splicing of transcripts encoding defense signaling proteins to dynamically increase immune receptor signaling capacity through deactivation of a key signal-buffering circuit.

## Main Text

Innate immunity has been described as a double-edged sword, providing essential protection in the face of attack, but detrimental to the host if allowed to persist^1^. Accordingly, immune responses are tightly regulated by complex layers of dynamic checks and balances. Rapid and reversible modulation of the signaling components requisite for this dynamic regulation commonly occurs through posttranscriptional, translational and posttranslational mechanisms without requiring *de novo* transcription^2–9^. Collectively these mechanisms rapidly modulate signaling pathways through stochiometric changes in the relative quantity or functional state of key regulators. However, elucidation of how these regulatory layers assemble into defined modules, particularly with respect to temporal dynamics and cause-effect relationships, remains challenging.

Pattern-recognition receptors (PRRs) recognize microbe-, herbivore-, and parasitic plant-associated molecular patterns to activate regulatory signaling networks that promote innate immune responses as a first line of defense^10, 11^. Following initiation of pattern-triggered immunity (PTI), endogenous plant hormones and messenger signals amplify initial inputs to coordinate immune outputs^11^. Among these, Plant Elicitor Peptides (Peps) have emerged as fundamental regulators of innate immunity across higher plants, with demonstrated ability to enhance resistance to a broad spectrum of insects, pathogens and nematodes in diverse species^12–18^. Peps are proteolytically released from PROPEP precursor proteins by the cysteine protease METACASPASE4 and activate PEP RECEPTORs (PEPRs) to coordinate downstream signals that trigger plant immune responses mediating resistance^12, 19–24^. Signaling by PEPRs requires SOMATIC EMBRYOGENESIS RECEPTOR KINASE (SERK) coreceptors and the plasma membrane-associated kinase BOTRYTIS INDUCED KINASE 1 (BIK1)^25–27^. Acting as a positive regulator of both reactive oxygen species- and WRKY transcription factor-mediated downstream responses, BIK1 is rate-limiting for signaling through pattern recognition receptor complexes, including PEPRs, and is continuously turned-over to maintain signaling homeostasis^28–30^. The E3 ligases PLANT U-BOX PROTEINs PUB25 and PUB26 facilitate turnover by ubiquitylating BIK1^31^. Phosphorylation of conserved residues in BIK1 and in PUB25/PUB26 by CALCIUM-DEPENDENT PROTEIN KINASE 28 (CPK28) enhances ubiquitin ligase activity and promotes BIK1 degradation. Through this mechanism, CPK28 negatively regulates immune receptor signaling by promoting BIK1 turnover^30, 31^.

Here we define a novel dynamic regulatory circuit acting on the CPK28 buffering system mediated by a newly-described RNA-binding protein, termed IMMUNOREGULATORY RNA-BINDING PROTEIN (IRR). In the absence of immune challenge, IRR physically interacts with CPK28 transcripts to promote canonical splicing into mRNA encoding full-length, functional proteins. Upon activation of PEPRs, IRR is transiently dephosphorylated, causing dissociation from *CPK28* transcripts. Disruption of IRR interaction with *CPK28* transcripts leads to increased levels of a retained-intron variant encoding a truncated protein that lacks EF-hand domains required for calcium-induced stimulation of kinase activity and exhibiting reduced functionality. Altered ratios of canonical versus retained-intron *CPK28* transcripts modulate PEPR signaling sensitivity, with proportional increases in the retained intron variant resulting in amplified immune signaling and defense. This dynamic circuit directly links PEPR-induced dephosphorylation of IRR with post-transcriptionally-mediated attenuation of CPK28 function, providing a rapid mechanism to modulate PEPR signaling capacity and immune outputs.

## Results

### Pep signaling promotes IRR dephosphorylation

To uncover post-translationally-modified regulators of immunity, we examined rapid Pep-induced protein phosphorylation changes in both Arabidopsis and maize. Suspension-cultured cells from each species were treated for 10 min with 100 nM *Arabidopsis thaliana* Pep1 (AtPep1) or *Zea mays* Pep3 (ZmPep3) respectively, and analyzed in comparison to water-treated controls using nano-liquid chromatography with tandem mass spectrometry (LC-MS/MS)^32^. In both species, a predicted RNA-binding protein containing an RNA Recognition Motif (RRM), IRR, was significantly dephosphorylated after Pep treatment (Supplementary Fig. 1), indicating a possible conserved role in Pep-induced signal transduction across plant species. Encoded by At3g23900 and GRMZM2G132936 genes in Arabidopsis and maize respectively, IRR contains zinc finger and RRM domains with a carboxyl-terminal region highly enriched in SR dipeptides (Fig. 1a).

**Figure 1.**
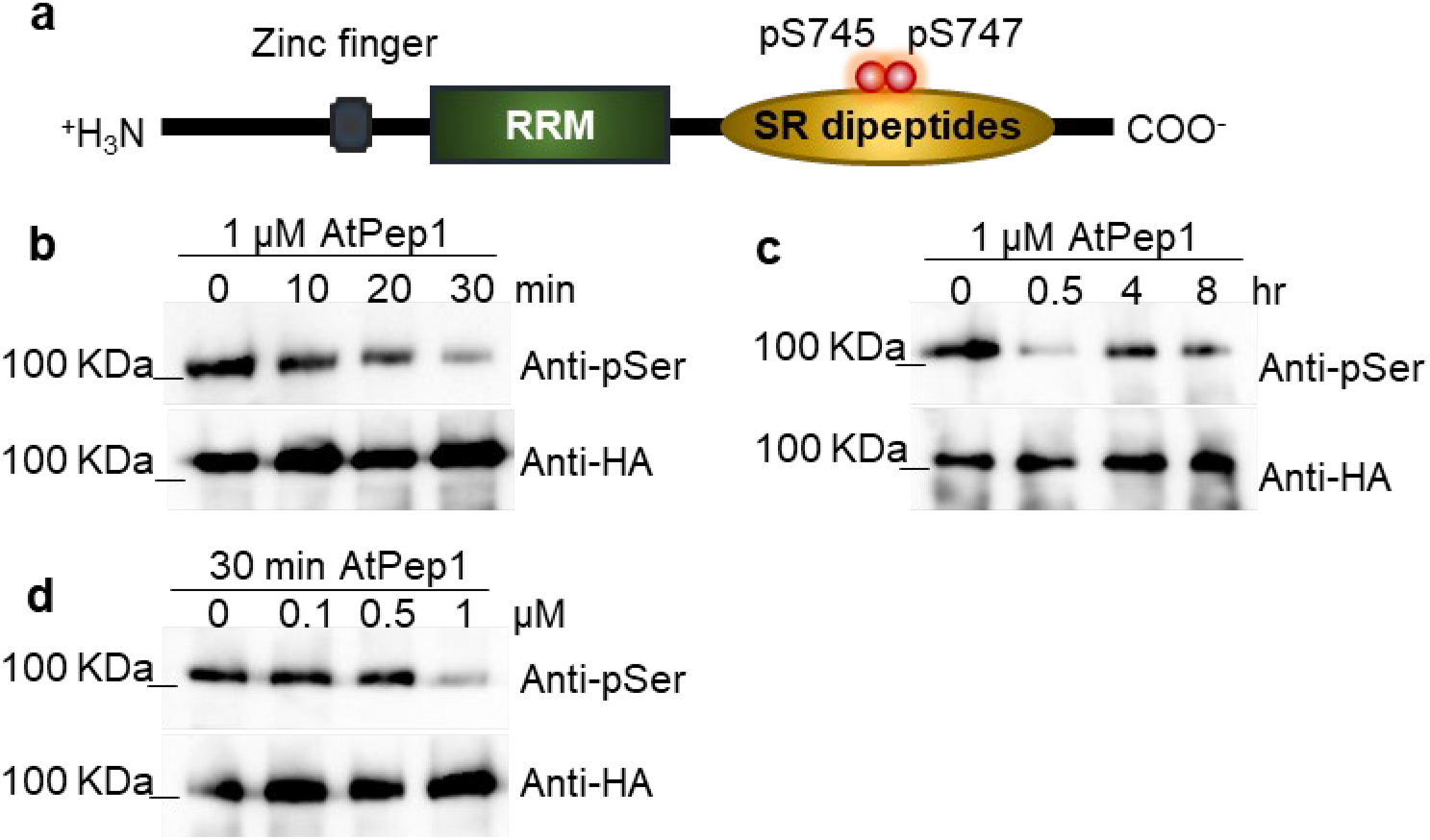
AtPep1 affects phosphorylation of IRR in a time- and concentration-dependent manner. **a**, IRR phosphorylation sites affected by AtPep1 treatment. **b**, p35S:IRR-3xHA plants were treated with 1 μM of AtPep1 for 0, 10, 20 and 30 min, and late time points, as 4 and 8 h (**c**). **d**, p35S:IRR-3xHA plants were treated with 0, 0,1, 0.5 and 1 μM AtPep1 for 30 min. IRR protein was subjected to immunoblot analysis with anti-phospho-Ser and anti-HA antibodies. Experiments in b–d were repeated three times independently, with similar results.

To confirm Pep-induced dephosphorylation of IRR *in planta*, Arabidopsis lines expressing a HA-tagged fusion of IRR (p35S:IRR-3xHA) were treated with AtPep1, followed by IRR immunoprecipitation and anti-phosphoserine antibody immunoblot analyses of phosphorylation states. Decreased IRR phosphorylation was observed within 30 min of AtPep1 treatment (Fig. 1b), while control anti-HA Western blots demonstrated no change in total IRR protein levels. AtPep1-induced dephosphorylation was transient, with a recovery to resting levels of IRR phosphorylation within 4 h (Fig. 1c). Decreased IRR phosphorylation also occurred in a concentration-dependent manner (Fig. 1d). To determine whether AtPep1-induced dephosphorylation was specific to IRR, phosphorylation of an unrelated serine/arginine (S/R)-rich RRM protein previously reported as a phosphoprotein^33^, termed SR45, was examined after Pep treatment using plants expressing an SR45-3xHA fusion protein (Supplementary Fig. 2). SR45 negatively regulates glucose and abscisic acid signaling during early seedling development, and recent transcriptional profiling studies have implicated it as a negative regulator of innate immunity^34, 35^. The phosphorylation levels of SR45 protein were unaltered by AtPep1 treatment (Supplementary Fig. 3), demonstrating that AtPep1-mediated dephosphorylation is not a general phenomenon in S/R-rich RRM proteins.

### Loss of function *irr* mutants display enhanced Pep-induced immune responses

To examine effects of IRR loss-of-function on AtPep1 response output, two T-DNA insertional knockout mutants of *IRR* (SALK_015201, termed *irr-1*, and SALK_066572, termed *irr-2*) were analyzed for altered sensitivity to AtPep1 treatment in comparison to an SR45 insertional mutant line (SALK_123442, *sr45-1*) as a control. Presence of T-DNA insertions and absence of target gene expression was confirmed for all lines by PCR (Supplementary Fig. 4b,c). The PEPR double knockout line, *pepr1/pepr2* (*pepr1/2*), which is fully insensitive to AtPeps, was employed as a negative control. As a component of defense activation, AtPep1 inhibits primary root elongation in Arabidopsis seedlings^22, 24^. Wild-type Col-0 (Wt) seedlings grown in media supplemented with 0.1 μM AtPep1 have primary roots approximately half the length of seedlings grown on water-supplemented medium (Fig. 2a). Consistent with AtPep hypersensitivity, primary roots of *irr1-1* and *irr1-2* were significantly shorter than Wt roots when grown on medium supplemented with both 0.1 μM and 1 μM AtPep1, indicating hypersensitivity to the peptide (Fig. 2a, Supplementary Fig. 5a), whereas *sr45-1* primary root growth was indistinguishable from Wt (Supplementary Fig. 5b,c). This result indicated a potential role for IRR as a negative regulator of AtPep1-induced responses. Additional layers of signaling and output downstream of AtPep1 were investigated, including production of second messengers, kinase activation and relative expression of Pep-associated marker genes. AtPep1 promotes generation of second-messenger reactive oxygen species (ROS) through stimulation of NADPH oxidase activity within min of PEPR activation^22^. While the timing of AtPep1-induced ROS production was the same in both Wt and *irr* knockouts, the magnitude of ROS generated was greater in *irr* lines (Fig. 2b, Supplementary Fig. 6a). AtPep1 treatment also stimulates phosphorylation-mediated activation of MAP KINASES (MPK) 3, 6 and 4, which are integral to many plant signaling pathways responsive to biotic and abiotic stresses^36, 37^. MPK3/6 activation after AtPep1 treatment was probed through Western blotting with anti-phospho-p44/42 MAPK antibody, revealing that phosphorylation of both MPK3 and MPK6 was more intense and prolonged in *irr* knockouts than in Wt, with detectable activity continuing 60 min after Pep treatment (Supplementary Fig. 6b). In correspondence with upregulation of second messenger and MAP kinase activities, expression of the AtPep1-responsive marker genes *PLANT DEFENSIN 1.2 (PDF1.2), TYROSINE AMINOTRANSFERASE 3 (TAT3*) and *PATHOGENESIS-RELATED PROTEIN 1* (*PR-1*) was also significantly increased 24 h after AtPep1 treatment in *irr* knockouts as compared to Wt (Fig. 2, c-e)^12^. The *irr-1* knockout phenotype was rescued through complementation with the IRR gene driven by its native promoter: two independent lines expressing pIRR:IRR-YFP in the *irr-1* background behaved as Wt plants in assays of AtPep1-induced root growth inhibition assay, ROS production and expression of the *PDF1.2* marker gene (Supplementary Fig. 7a-c).

**Figure 2.**
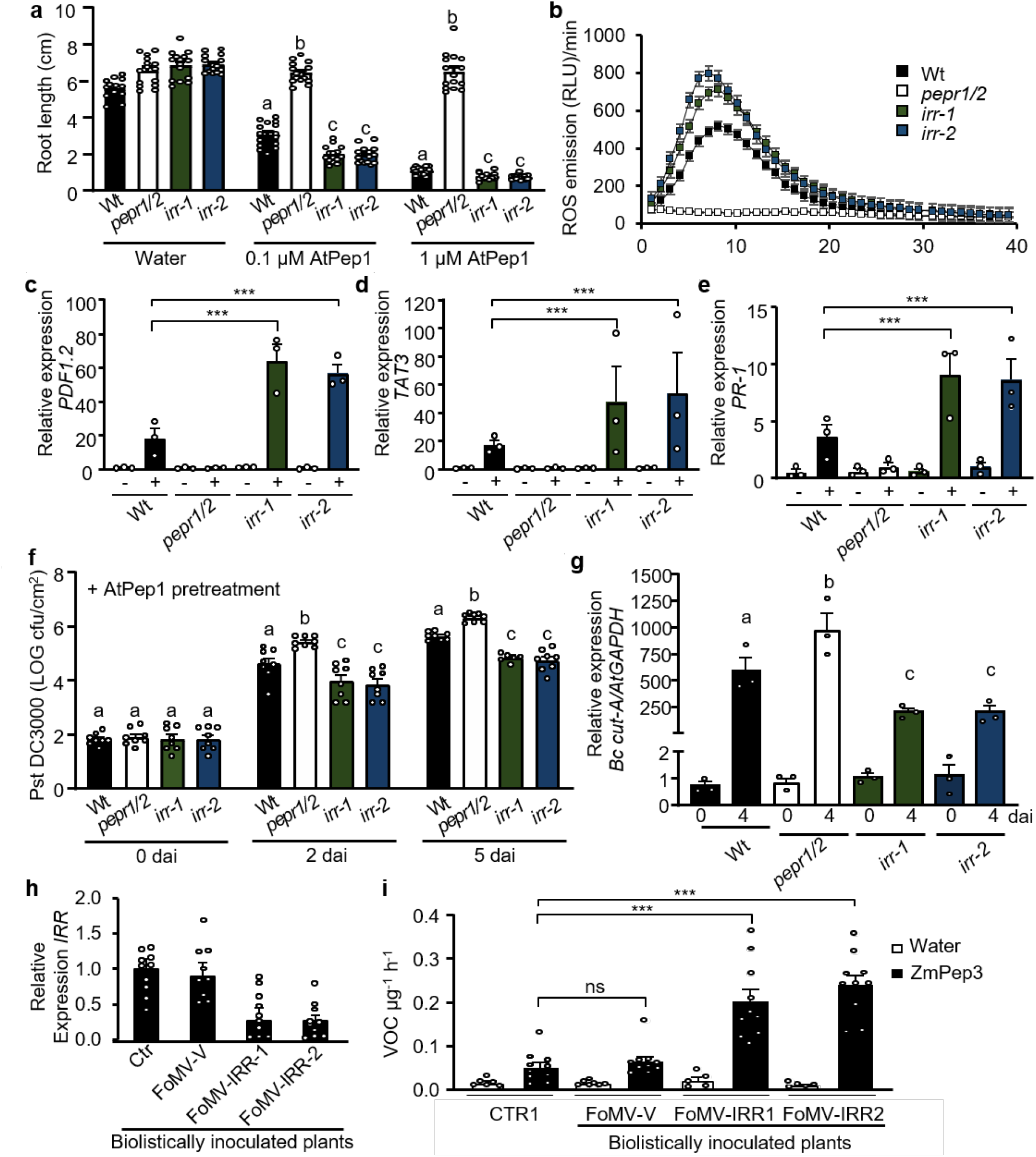
*IRR* mutants are hypersensitive to Pep treatment. **a**, Arabidopsis seedlings were treated with AtPep1 (0.1 μM and 1 μM) or water, and the root length was measured after 15 days of treatment. The values are the mean ± SD of 15 seedlings. The means with the same letter are not significantly different from each other (Tukey’s test, P <0.01). **b**, ROS production was registered continuously with cut and pre-incubated leaf pieces using the luminol bioassay and monitored during 40 min after addition of 1 μM AtPep1. **c**, The relative expression level of AtPep1-inducible genes *PDF1.2, TAT3*, (**d**) and *PR-1* (**e**) was determined by real-time qRT-PCR. mRNA from entire seedlings treated with water (-) and 1 μM AtPep1 (+) for 24 h was analyzed. *GAPDH* was used as reference gene. Error bars indicate the SD of three biological replicates. **f**, Pst DC3000 infection assay of wild-type, *pepr1/2, irr-1* and *irr-2* plants after 1 μM AtPep1 pretreatment. The peptide was infiltrated 48 h prior to infection. Bars indicate samples just after inoculation (0), 2 and 5 days after inoculation (dai). Error bars indicate SD. n=8. **g**, *Botrytis cinerea* infection in wild-type, *pepr1/2* and *irr* mutants. Quantification of in planta growth of *B. cinerea*. qPCR was used to analyze the relative genomic DNA level of *B. cinerea Cutinase A (Bc cut-A*) compared with Arabidopsis *GAPDH (Bc cut-A/AtGAPDH*). Error bars indicate SD. n=15. Columns with the same letter are not significantly different (Tukey test. P ≤ 0.01). Experiments in a–g were repeated at least three times independently, with similar results. **h**, *IRR* knockdown maize plants analysis. *IRR* gene (GRMZM2G132936) analysis expression. The bar graphs display the relative expression levels of *IRR* mRNA in leaf samples determined by real-time qRT-PCR. mRNA from leaf 5 was analyzed. Expression of *RPL17* gene was used as reference gene. **i**, Total volatile organic compounds (VOCs) from maize leaves 16 h posttreatment with water or with 5 μM ZmPep3. Leaf 5 was used. All error bars indicate the SD of 5-10 biological samples. Single and triple asterisks indicate P <0.001 and P<0.05 (Student’s t-test), respectively; ns, not significant. Experiments in h,i were repeated two times independently, with similar results.

To assess whether enhanced AtPep1-induced immune responses in *irr* lines translated to increased disease resistance, plants were challenged with both the hemibiotrophic bacterial pathogen, *Pseudomonas syringae* pv tomato DC3000 (Pst DC3000), and the necrotrophic fungal pathogen, *Botrytis cinerea*. When inoculated with Pst DC3000 by leaf infiltration, bacterial proliferation was similar in both wild-type and *irr-1* plants after two and five days (Supplementary Fig. 8a), whereas the *pepr1/2* double mutant showed more susceptibility to Pst DC3000, as previously reported^24^. In contrast, after priming immunity through pre-infiltration of leaves with AtPep1 48 h prior to Pst DC3000 inoculation, bacterial proliferation was significantly reduced in *irr-1* and *irr-2* knockout plants when compared to wild-type (Fig. 2f). Upon challenge with *B. cinerea*, four days post-inoculation, the average lesion area of *irr* leaves was smaller than for Wt (Supplementary Fig. 8b). Correspondingly, quantification of fungal proliferation through quantitative RT-PCR analysis of relative ratio of *B. cinerea Cutinase A* DNA versus *A. thaliana GAPDH* revealed significantly lower pathogen levels in both *irr* mutant lines as compared to wild-type (Fig. 2g).

To investigate IRR function in maize, a Viral-Induced Gene Silencing (VIGS) system derived from Foxtail Mosaic Virus (FoMV) was deployed to silence maize *IRR* (Supplementary Fig. 9a,b)^38^. Maize var. B73 seedlings were biolistically inoculated with FoMV-derived vectors carrying two different IRR sequence fragments, designated as FoMV-IRR-1 and FoMV-IRR-2. Control plants were inoculated with FoMV vector carrying no insert (FoMV-V). For each FoMV construct, 10 plants confirmed as infected were selected for further analysis. To evaluate relative *IRR* silencing, *IRR* transcript levels were compared among leaves of plants infected with FoMV-IRR-1/2, empty vector FoMV-V and uninoculated control plants using qRT-PCR. Expression levels of *IRR* were similar in uninoculated and FoMV-V-infected maize plants, demonstrating that FoMV-V infection alone did not significantly affect *IRR* gene expression (Fig. 2h). However, infection with either of the FoMV-IRR constructs significantly reduced *IRR* expression (Fig. 2h). As *irr* knockout Arabidopsis mutants are hypersensitive to AtPep1, maize FoMV *IRR* knockdown plants were tested for sensitivity to ZmPep3. In maize, ZmPep3 is a potent inducer of herbivore-associated volatile organic compounds (VOC) that serve as indirect defenses by recruiting parasitic wasps to attack Lepidopteran herbivore pests feeding on leaves^14^. VOC emission from FoMV-infected maize leaves after ZmPep3 treatment was measured by gas chromatography, and total VOCs emitted after peptide treatment was significantly higher from maize plants inoculated with FoMV-IRR-1/2 than from control plants (Fig. 2i). A direct comparison of *IRR* mRNA transcript levels versus VOC emission levels in FoMV-IRR-1/2 confirmed that lower *IRR* expression correlated with higher VOC emission (Supplementary Fig. 9c,d). Together these experiments support IRR function as a conserved negative regulator of Pep-induced immune responses.

### *irr* knockouts exhibit broad changes in defense gene expression and splicing patterns

To better understand potential mechanisms underlying *irr* hypersensitivity to Pep treatment, global transcriptional patterns in *irr-1* and wild-type plants were profiled by RNA-seq 24 h post-treatment with either water or 1 μM AtPep1. Untreated *irr-1* plants demonstrated extensive dysregulation of genes relating to the immune response, with over 600 genes differentially regulated in *irr-1* compared to wild-type (Supplementary Fig. 10, Supplementary Table 1). Analysis of Gene Ontology (GO) for transcripts with increased basal expression levels in *irr-1* revealed significant enrichment of immunity-related terms, with top categories including response to stimulus, defense response, immune system process and programmed cell death (Fig. 3a, Supplementary Table 2).

**Figure 3.**
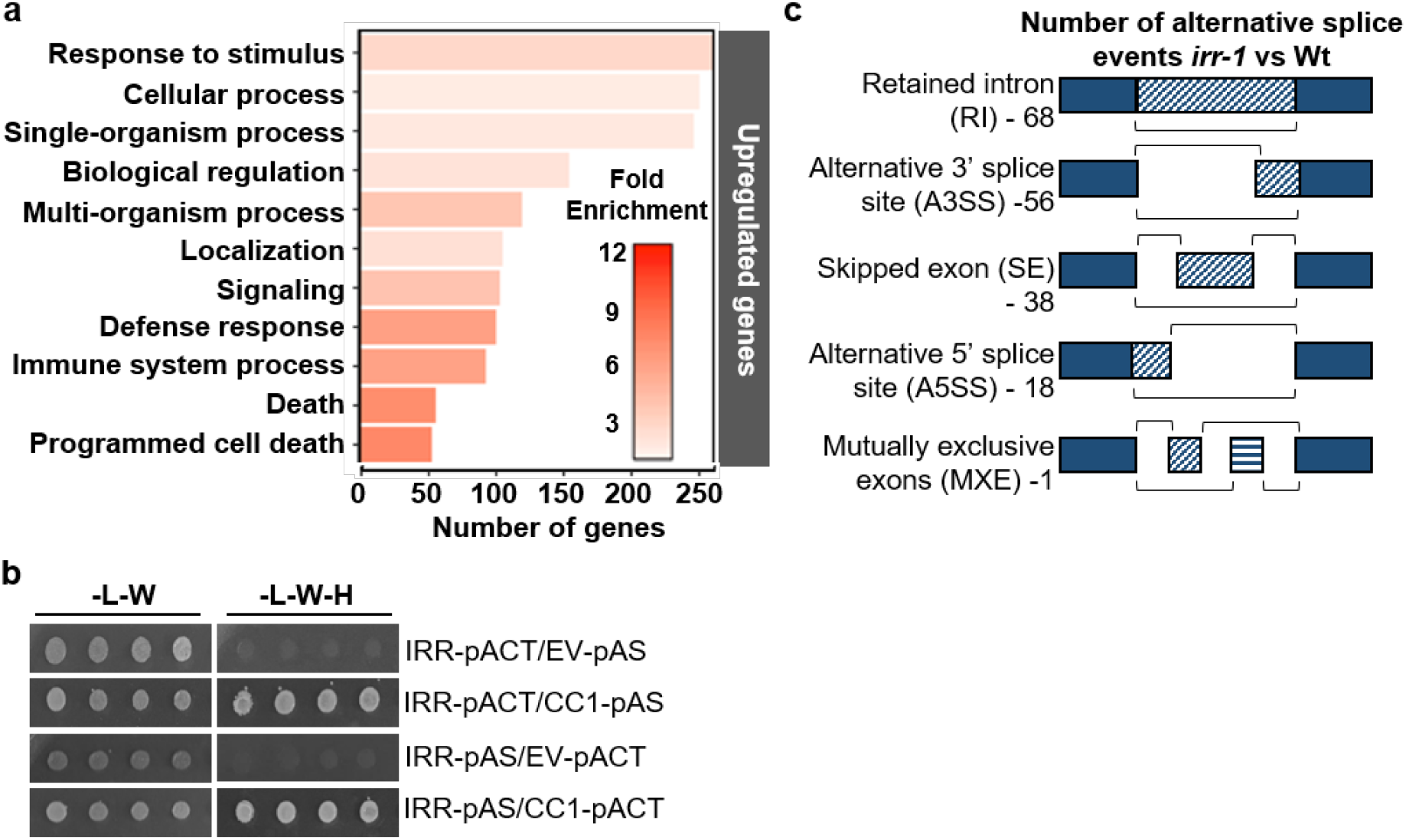
IRR is implicated in defense and alternative splicing. **a**, GO term distribution of upregulated expressed genes in *irr-1* versus Wt. **b**, IRR interacts with CC1-splicing factor in yeast. Yeast strain AH109 was co-transformed with IRR and CC1-splicing factor encoding proteins. The transformants were selected in media lacking leucine and tryptophan (-L-W), and interaction was tested in media lacking leucine, tryptophan and histidine (-L-W-H). This experiment was performed three times with similar results. c, Number of alternative splice events in *irr-1* vs Wt. For all experiments, three biological replicates were analyzed.

RNA-binding proteins with serine/arginine-enrichment and RRM domains are established as key to precursor-mRNA processing and alternative splicing in eukaryotes ^39, 40^, thus we considered related roles for IRR. Notably, widespread changes in splicing patterns were observed in *irr-1* as compared to wild-type, with differences in retained-intron, alternative 3’ splice site, skipped-exon, alternative 5’ splice site and mutually exclusive exon events (Fig. 3b, Supplementary Table 3). Among the alternative splicing patterns observed in *irr-1* knockout, retained-intron events were most abundant. Numerous transcripts encoding defense signaling proteins exhibited differing ratios of retained-intron variants encoding premature stop codons that would predictably result in truncated proteins and potentially modified functions (Supplementary Fig. 11). Negative regulators of plant immune signaling, such as *CPK28*, LESION-SIMULATING DISEASE 1 (*LSD1*), and JASMONATE-ZIM-DOMAIN PROTEIN 4 (*JAZ4*) all exhibited increased levels of retained-intron transcripts, whereas a positive regulator of immunity, Cysteine-rich Receptor-like Kinase 13 (*CRK13*) had markedly decreased levels of a retained-intron isoform ^30, 41–44^ (Supplementary Fig. 12a). To validate occurrence of retained-intron (RI) events observed through RNA-seq analysis, qRT-PCR was performed with primer sets unique to the intron of target genes that was retained at different ratios in *irr-1* and wild-type plants. Amplification of the RI variant was performed in parallel with amplification using primers for the canonical splice form so that relative ratios could be compared. These analyses confirmed that in the *irr-1* mutant, the ratio of *CPK28, LSD1* and *JAZ4* RI splice variants increased compared to wild-type, whereas the ratio of *CRK13* RI splice variant decreased (Supplementary Fig. 12b).

In support of a role in mRNA splicing, IRR has been predicted to physically interact with the CC1-LIKE SPLICING FACTOR encoded by At2g16940^45^. Using a yeast two-hybrid system with co-expression of IRR and CC1 fused to bipartite transcription factor halves, IRR was found to physically interact with CC1 as predicted (Fig. 3c). Co-expression of CC1 with SR45, which has been demonstrated to mediate pre-mRNA splicing, also yielded a positive interaction^46^ (Supplementary Fig. 13). Because *sr45-1* knockout does not share the AtPep1-hypersensitive phenotype of *irr* knockouts, we conclude that while CC1 interaction with these proteins likely facilitates function, it is unlikely to contribute to target specificity.

### The *CPK28*-RI splice variant underlies reduced negative regulation

Retained-intron variants of transcripts encoding CPK28 were among the most significantly increased alternative splicing events observed in *irr-1* knockouts, with a ratio of *CPK28*-RI to total *CPK28* transcripts six-fold higher than in Wt plants (Fig. 4a). Given the known function of CPK28 negatively regulating AtPep1-PEPR complex activity by promoting turnover of the rate-limiting component BIK1, the role of IRR-mediated changes in *CPK28* transcript splicing was examined^30, 31^. Calcium-dependent protein kinases are most commonly activated through exposure of the catalytic site as a result of conformational changes triggered by calcium ion-binding to the four regulatory EF hand domains^47, 48^. Partial losses of these EF hand domains can inhibit Ca^2+^-induced conformational changes, resulting in inactive CPKs with shielded catalytic domains^47^. The truncated CPK28 protein encoded by the *CPK28*-RI splice variant lacks two C-terminal EF hands (Fig. 4b), and was predicted to exhibit compromised catalytic activity as compared to full-length CPK28. Production of truncated CPK28 from *CPK28*-RI was confirmed through expression of cDNA encoding either canonical *CPK28* or *CPK28*-RI as both a YFP-fusion in Arabidopsis and a GST-fusion in *E. coli*. In both cases, *CPK28*-RI yielded a truncated fusion protein of the expected size (Fig 4c,d). Using *in vitro* kinase activity assays, GST-CPK28 actively auto-phosphorylated and trans-phosphorylated both GST-BIK1 protein and the catalytically inactive variant GST-^BIK1K105A/K106A 27, 30^, while GST-CPK28-RI had greatly reduced phosphorylation activity (Fig. 4d).

**Figure 4.**
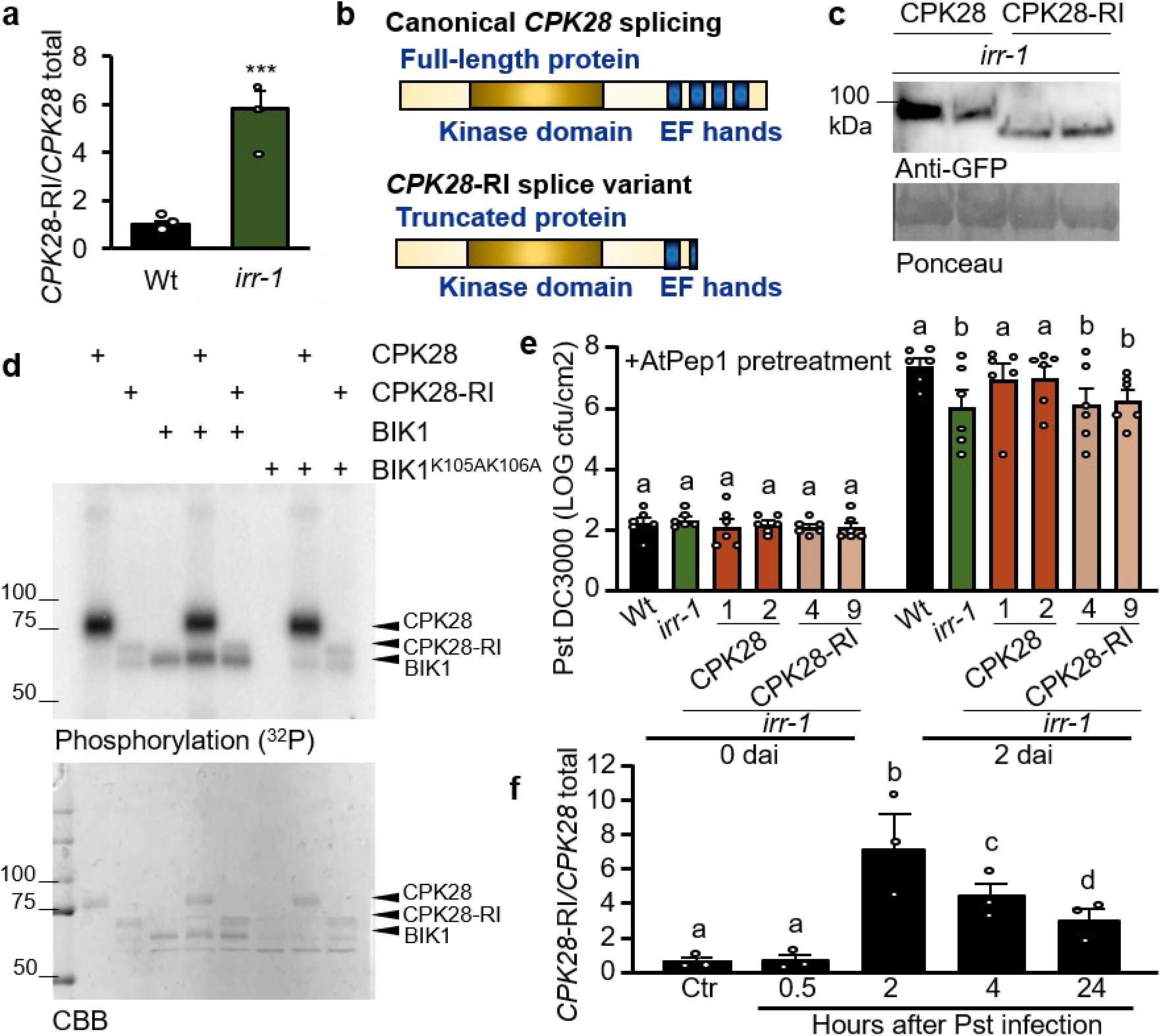
IRR affects the ratio of *CPK28* retained-intron splice variants and CPK28 function. **a**, Relative expression of *CPK28*-RI splice variant in wild-type versus *irr-1* plants. **b**, Canonical splicing of the CPK28 transcript from a full-length protein containing four EF hands versus the retained intron-CPK28 splice variant results in a truncated protein missing EF hands. **c**, The proteins CPK28 and CPK28-RI were detected by Western blot. Proteins were extracted from leaves of transgenic plants expressing pCPK28-CPK28-YFP and pCPK28-CPK28-RI-YFP in *irr-1* background. Anti-GFP antibody was used to detect both proteins. Two independent events per line were analyzed. CPK28-YFP, 85 kDa; CPK28-RI-YFP, 75 kDa. Ponceau was used for verification of protein loading. d, Autoradiograph showing incorporation of radioactive phosphate in the GST-fused recombinant proteins following kinase assay *in vitro*. CBB stains are included as controls. **e**, Pst DC3000 infection assay of wild-type, *irr-1*, irr-1/pCPK28:CPK28-YFP and *irr*-1/pCPK28:CPK28-RI-YFP plants after 1 μM AtPep1 pre-treatment. The peptide was infiltrated 48 h prior to infection. Bars indicate samples 0 and 2 days after inoculation (dai). Error bars indicate SD. n=6. **f**, Relative expression of the *CPK28*-RI splice variant transcripts in wild-type plants after Pst DC3000 infection, analyzed by qRT-PCR. Plants were infected with Pst DC3000, and the analysis was performed 0.5, 2, 4 and 24 h after infection. Non-infected plants were used as control (Ctr). Triple asterisks indicate P< 0.001 (Student’s t-test). Columns with the same letter are not significantly different (Tukey test. P ≤ 0.01). For experiments a and f, three biological replicates were analyzed. Experiments were repeated at least two times independently, with similar results.

We hypothesized that a mechanism underlying *irr-1* hypersensitivity to AtPep1 treatment is reduced negative regulation of PEPR signaling by CPK28 due to a loss of IRR function in promoting canonical splicing of *CPK28* transcript to produce a functional full-length protein. In this case, increased expression of the canonical CPK28 transcript should fully or partially rescue the *irr-1* phenotype. To test this hypothesis, Arabidopsis plants were transformed with cDNA constructs encoding both transcript variants in the *irr-1* mutant background under control of the CPK28 native promoter and fused to a YFP tag. Two independent events for each transformation were selected, and protein expression confirmed by Western blot, with plants expressing pCPK28:CPK28-RI-YFP producing a smaller protein than plants expressing pCPK28:CPK28-YFP (Fig. 4c). AtPep1-induced resistance to *Pseudomonas syringae* pv. tomato DC3000 (Pst) was examined in *irr-1* plants expressing pCPK28:CPK28-YFP versus pCPK28:CPK28-RI-YFP. As previously observed, when pretreated with AtPep1 48 h prior to inoculation, *irr-1* knockout plants were more resistant to Pst infection than Wt as measured by decreased bacterial proliferation after two days. In *irr-1* plants expressing pCPK28:CPK28-YFP, Pst proliferation was similar to Wt, confirming that increased levels of canonical *CPK28* transcript were able to rescue the *irr-1* phenotype (Fig. 4e). In contrast, expression of pCPK28:CPK28-RI-YFP in *irr-1* did not rescue the AtPep1-hypersensitive phenotype, as these lines exhibited the same enhanced AtPep1-induced restriction of bacterial proliferation as *irr-1*. Together the evidence indicates that: (1) truncated CPK28-RI protein is not fully functional as a negative regulator, and (2) the inability of *irr* knockout plants to promote canonical *CPK28* splicing for production of a functional full-length protein that negatively regulates PEPR signaling is a significant contributor of *irr* hypersensitivity to AtPep1. Furthermore, relief of CPK28-mediated negative regulation of immunoregulatory receptor complex signaling appears integral to plant defense responses: Pst infection triggers an increased proportion of *CPK28*-RI to total *CPK28* transcripts (Fig. 4f), demonstrating that *CPK28* alternative splicing is a common mechanism to dynamically enhance plant immunity.

### Accumulation of *CPK28*-RI transcript is dependent on IRR phosphorylation state

To ascertain whether levels of the *CPK28*-RI splice variant were affected by Pep signaling, Wt plants were treated with AtPep1 and expression of total *CPK28* and *CPK28*-RI transcripts analyzed by qRT-PCR. Notably, AtPep1 promoted significant increases in proportional *CPK28*-RI levels at 30 min (Fig. 5a), temporally coinciding with peak AtPep1-induced dephosphorylation of IRR (Fig. 1c). Elevated proportions of *CPK28*-RI returned to initial levels after 4 h (Fig. 5a), concurrent with recovery of IRR phosphorylation post-AtPep1 treatment. To determine whether IRR phosphorylation state contributed to *CPK28*-RI accumulation, *irr-1* plants overexpressing an HA-tagged fusion of the phospho-abolishing mutant of IRR, IRR^S745A,S747A^, were examined. The *irr*-1:IRR^S745A,S747A^-3xHA plants exhibited higher levels of *CPK28*-RI than Wt or *irr-1*:IRR-3xHA plants, demonstrating a phenotype equivalent to *irr-1* knockout plants and to *irr-1* plants carrying the empty vector, *irr-1*:3xHA (Fig. 5b). The inability of IRR^S745A,S747A^ to reduce proportional *CPK28*-RI levels in the *irr-1* background supports dephosphorylation of IRR as a contributor to accumulation of *CPK28*-RI variant transcripts after AtPep1 treatment. Similarly, IRR^S745A,S747A^ failed to rescue the AtPep1-hypersensitive phenotype of *irr-1*. Expression of the AtPep1-induced marker gene *PDF1.2* remained elevated in *irr-1:* IRR^S745A,S747A^-3xHA lines, whereas *irr-1* lines complemented with IRR phenocopied Wt (Fig. 5c). Furthermore, *irr-1*:IRR^S745A,S747A^-3xHA plants exhibited enhanced AtPep1-induced restriction of Pst proliferation comparable to *irr-1*, whereas Pst proliferation in *irr-1*:IRR-3xHA plants was similar to Wt (Fig. 5d). Together these results suggest that phosphorylated IRR acts as a negative regulator of AtPep1-induced immune responses. In unchallenged wild-type plants, the population of IRR protein is predominantly phosphorylated on pS745 and pS747, and active as a negative regulator. However, upon perception of AtPep1, transient dephosphorylation of IRR temporarily attenuates negative regulatory function. When IRR is absent, as in *irr* knockouts, IRR-mediated negative regulation is fully relieved and results in enhanced immune responses.

**Figure 5.**
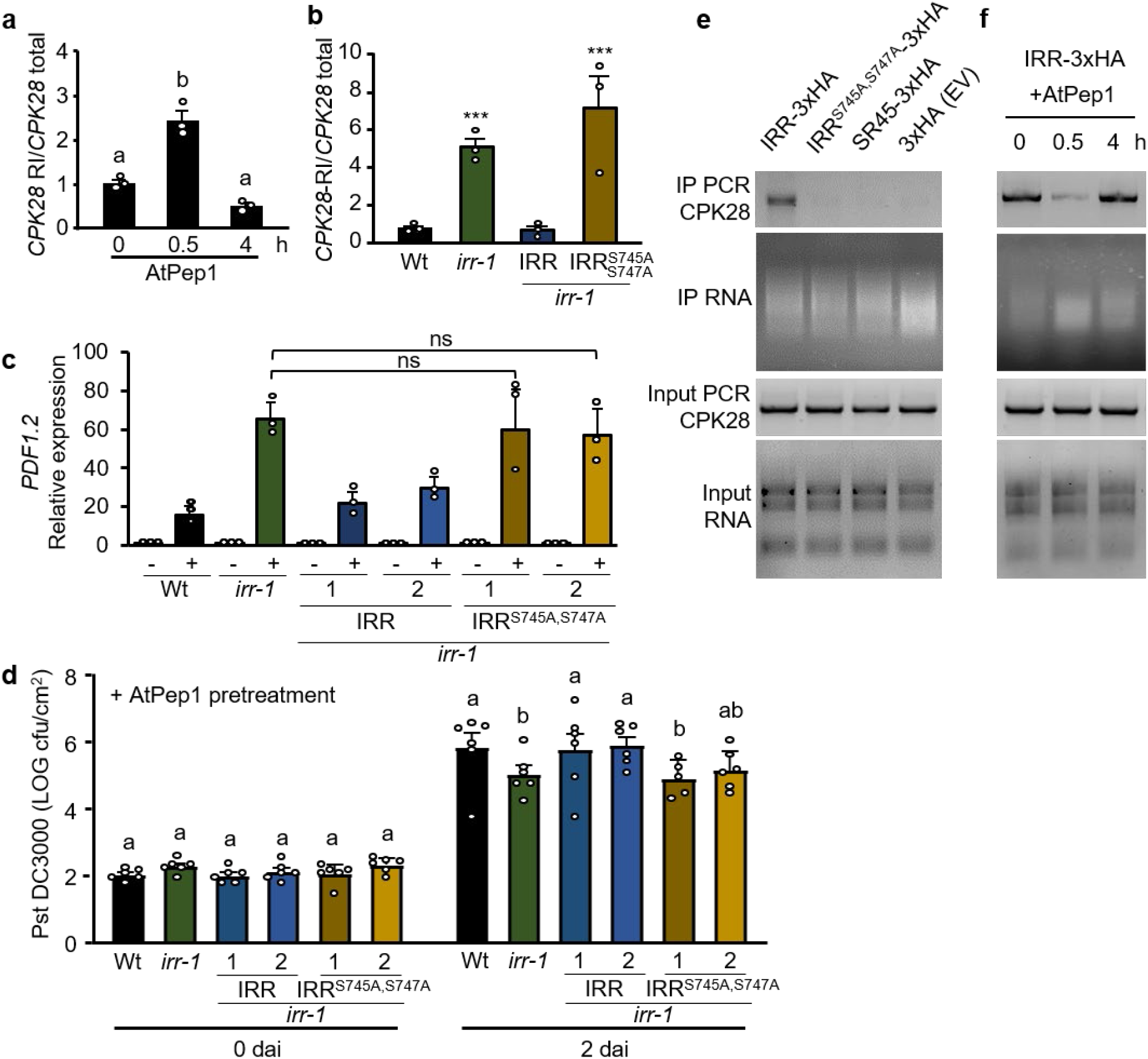
IRR interacts with *CPK28* transcript in a phosphorylation-dependent manner. **a**, Relative expression of *CPK28*-RI splice variant in transgenic plants overexpressing IRR-3xHA, analyzed by qRT-PCR. Plants p35:IRR-3xHA were treated with 1 μM AtPep1 for 0, 30 min and 4 h. **b**, Relative expression of *CPK28* RI in Wt, *irr-1* and transgenic lines overexpressing triple HA-tagged IRR (*irr-1*:IRR) and IRR^S745A,S747A^ (*irr-1*: IRR^S745A,S747A^). *ACTIN2* was used as refence gene. **c**, The relative expression of AtPep1-inducible gene *PDF1.2* was determined by real-time qRT-PCR. mRNA from entire seedlings treated with water (-) or 1 μM AtPep1 (+) for 24 h was analyzed. Two independent events per transgenic line overexpressing triple HA-tagged IRR (*irr-1*:IRR) and IRR^S745A,S747A^ (*irr-1*:IRR^S745A,S747A^) were used. *ACTIN2* was used as reference gene. Error bars indicate the SD of three biological replicates. **d**, Pst DC3000 infection assay of wild-type, *irr-1* and *irr-1* overexpressing triple HA-tagged IRR (*irr-1*: IRR) and IRR^S745A,S747A^ (*irr-1*:IRR^S745A,S747A^) plants after 1 μM AtPep1 pre-treatment. The peptide was infiltrated 48 h prior to infection. Bars indicate samples 0 and 2 days after inoculation (dai). **e**, RNA immunoprecipitated (RIP) from transgenic plants overexpressing IRR-3xHA, IRR^S745A,S747A^-3xHA, SR45-3xHA and 3xHA (Empty vector, EV). RIP-PCR was performed to detect *CPK28* transcripts. **f**, RIP from transgenic plants overexpressing IRR-3xHA after 1 μM AtPep1 treatment for 0, 30 min and 4 h. RIP-PCR was performed to detect *CPK28* transcripts. The protein-RNA complex was immunoprecipitated using anti-HA magnetic beads. Input RNA was extracted from all samples as control. PCR reaction using primers to detect *CPK28* transcript using as a template cDNA synthetized from the input RNAs. Error bars indicate SD. n=6. Triple asterisks indicate P< 0.001 (Student’s t-test). Columns with the same letter are not significantly different (Tukey test. P ≤ 0.01). Experiments were repeated three times independently, with similar results.

### IRR physically interacts with *CPK28* transcript in a phosphorylation-dependent manner

To better understand how dephosphorylation of IRR might contribute to function, behavior of the IRR^S745A,S747A^ mutant protein relative to wild-type IRR was assessed. Subcellular localization of YFP-tagged IRR^S745A,S747A^ was examined as compared to IRR-YFP, and found to be unaffected (Supplementary Fig. 14). For many serine/arginine-rich RNA-binding proteins that function in precursor-mRNA processing, phosphorylation promotes interaction with other proteins in the splicing complex^49^, so the effect of mutant IRR^S745A,S747A^ on physical interactions with the CC1 splicing factor was also examined. As determined by plate-based yeast two-hybrid and monitoring of yeast growth dynamic in liquid medium, mutation of the phosphorylation sites to alanine does not impair interaction with the CC1-splicing factor, nor alter interaction affinity (Supplementary Fig. 15a-c). To investigate whether AtPep1-induced dephosphorylation affects IRR stability and turnover, *irr-1* plants overexpressing triple HA-tagged IRR were treated with AtPep1. No change in total IRR protein levels was observed by Western blot after AtPep1 treatment (Supplementary Fig. 16a). However, cotreatment with AtPep1 and the proteasome inhibitor MG132, or the protein translation inhibitor cycloheximide (CHX), revealed that phosphorylation state may affect turnover rate. Treatment with MG132 or CHX alone did not affect IRR levels, but with cotreatments, IRR accumulated 30 and 120 min after MG132/AtPep1 treatment, and was depleted after CHX/AtPep1 treatment (Supplementary Fig. 16a,b). Mutant IRR^S745A,S747A^ protein levels were significantly decreased by CHX treatment in both the absence and presence of AtPep1, suggesting that dephosphorylation of IRR promotes protein turnover (Supplementary Fig. 16c).

To ascertain whether IRR regulation of *CPK28* transcript splicing might occur through direct physical interaction, RNA immunoprecipitation (RIP) coupled with PCR analysis was used to test for IRR-CPK28 transcript interactions. RIP-PCR demonstrated that IRR physically interacts with *CPK28* mRNA, with a *CPK28* fragment amplified from RNA co-immunoprecipitating with IRR (Fig. 5e). The RNA-binding protein SR45 was used as a negative control since SR45-interacting transcripts have previously been identified by RIP-Seq, and *CPK28* is not among them^50^. As expected, amplification of RNA co-immunoprecipitated with SR45 did not yield a CPK28 fragment, nor did amplification of RNA co-immunoprecipitated with an unfused triple-HA tag (Fig. 5e). Because mutant IRR^S745A,S747A^ fails to promote canonical splicing of *CPK28*, in contrast to wild-type IRR, dependence of the IRR-CPK28 interaction on pSer745 and pSer747 was probed. No *CPK28* fragment was amplified from RNA co-immunoprecipitation with IRR^S745A,S747A^ (Fig. 5e), indicating that abolished phosphorylation at these sites disrupts physical interaction of IRR with *CPK28* transcript in addition to eliminating IRR-stimulated canonical splicing of *CPK28*. All proteins were detected after protein-RNA complex immunoprecipitation (Supplementary Fig. 17).

Because phospho-abolishing mutations of IRR also abolished interaction of IRR with *CPK28* transcript, the effects of AtPep1-mediated transient dephosphorylation of IRR on *CPK28* transcript interactions were investigated. Coimmunoprecipitation of triple HA-tagged IRR with associated RNA was performed 0, 0.5, or 4 h post-treatment with AtPep1 to compare with IRR dephosphorylation dynamics (Fig. 1a-b). *CPK28* transcript was abundant in IRR-interacting RNA at 0 h, but declined in RNA coimmunoprecipitated with IRR 30 min after AtPep1 treatment (Fig. 5f). Thus AtPep1-induced dephosphorylation of IRR parallels a physically disrupted interaction with *CPK28* transcript. Predictably, 4 h after AtPep1 treatment, when IRR phosphorylation levels have recovered (Fig. 1b), increased amplification of *CPK28* transcript from coimmunoprecipitated RNA was observed, indicating a reestablishment of IRR/CPK28 interaction (Fig. 5f). For all samples, input and coimmunoprecipitated RNA was determined to be of similar quantities and *CPK28* transcript amplified similarly from input RNA collected from samples prior to coimmunoprecipitation (Fig. 5e,f).

## Discussion

Dynamic plant immunity is essential to effectively resisting biotic attack^3–7^. We have identified a new RNA-binding protein, IRR, conserved in both maize and Arabidopsis that acts as a rapid molecular switch to impact response strength. In addition to identifying alternative splicing events mediated by IRR, we have demonstrated how dynamic and site-specific post-translational modification of IRR regulates physical interaction with, and alternative splicing of, transcripts encoding the key defense regulator CPK28 to affect immune response outputs. We further demonstrate that targeted mutation of IRR can confer enhanced disease resistance.

Based on our findings, we propose a model by which IRR dynamically regulates the CPK28 immunomodulatory buffering system (Fig. 6). Prior to immune challenge, IRR is predominantly phosphorylated at S745 and S747, and physically interacts with *CPK28* transcripts, facilitating canonical splicing to produce a full-length, functional protein (Fig. 6a). Full-length CPK28 promotes BIK1 turnover to suppress immune receptor signaling by phosphorylating both BIK1 and the ubiquitin ligases PUB25 and PUB26 to promote BIK1 ubiquitylation and degradation^30, 31^. Upon Pep-induced activation of PEPRs, IRR is transiently dephosphorylated and dissociates from *CPK28* transcripts, resulting in increased levels of the retained-intron *CPK28*-RI variant (Fig. 6b). The truncated protein encoded by *CPK28*-RI lacks EF hand motifs required for calcium-induced stimulation of kinase activity. Increased levels of this less-active CPK28-RI protein attenuate CPK28-mediated BIK1 degradation and temporarily enhance signaling capacity of PEPR complexes to amplify defense outputs. Re-phosphorylation of IRR facilitates equilibration back to immunoregulatory homeostasis, with CPK28 again buffering receptor complex function (Fig. 6c). Together this study defines a dynamic circuit that directly links PEPR-induced dephosphorylation of IRR with post-transcriptionally mediated attenuation of CPK28 function, revealing a new mechanism to modulate PEPR signaling capacity and immune response outputs.

**Figure 6.**
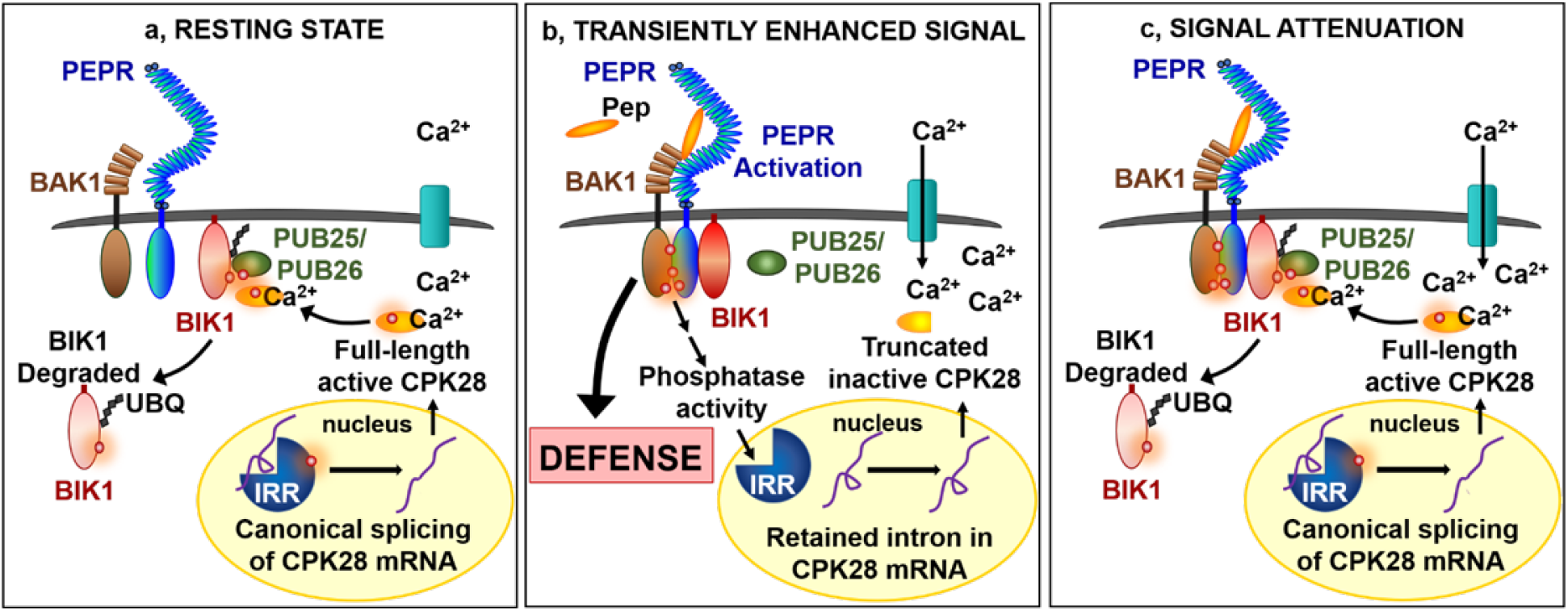
Model for IRR dynamic regulation of the CPK28 immunomodulatory buffering system. **a**, <u>Resting state</u>: phosphorylated IRR interacts with transcripts encoding *CPK28*, facilitating canonical splicing into mRNA that produce a full-length protein which functions as a negative regulator of immune receptor complex signaling by phosphorylating PUB25/26 ubiquitin ligases to promote ubiquitylation and subsequent degradation of BIK1. **b**, <u>Transiently enhanced signal</u>: AtPep1 activates PEPRs-BAK1 coreceptor complex, IRR is transiently dephosphorylated and dissociates from *CPK28* transcripts, resulting in temporary accumulation of a retained intron variant encoding a premature stop codon that yields a truncated and inactive CPK28 protein variant. This results in increased PEPR signaling capacity and enhanced immune output. **c**, <u>Signal attenuation</u>: recovery of IRR phosphorylation promotes canonical CPK28 splicing, reconstituting CPK28 buffering of the receptor complex through BIK1 turnover.

## Acknowledgments

The authors thank Prof. Steven A. Whitham (Iowa State University Plant Sciences Institute) for providing the constructs for VIGS experiments.

## Author contributions

A.H. and K.D. conceived the project. K.D. conducted experiments. A.H. and K.D. analyzed data and wrote the manuscript. P.W. performed MAP kinase assay, confocal microscopy experiments and phylogenetic analysis. E.P. analyzed leaf volatile emission and helped to edit figures. Y.T. performed in-gel kinase activity assays. C.V. assisted with generation of transgenic plant lines. Z.S. and S.P.B. performed phosphoproteomic analysis. J.I.S. contributed critical experimental resources.

## References

1. Kobayashi, KS and Flavell, RA, Shielding the double-edged sword: negative regulation of the innate immune system. J Leukoc Biol, 2004. 75(3): p. 428–33.

2. Gassmann, W, Alternative splicing in plant defense. Curr Top Microbiol Immunol, 2008. 326: p. 219–33.

3. Liu, J, Qian, C, and Cao, X, Post-Translational Modification Control of Innate Immunity. Immunity, 2016. 45(1): p. 15–30.

4. Xu, G, Greene, GH, Yoo, H, Liu, L, Marques, J, Motley, J, and Dong, X, Global translational reprogramming is a fundamental layer of immune regulation in plants. Nature, 2017. 545(7655): p. 487–490.

5. Nuhse, TS, Bottrill, AR, Jones, AM, and Peck, SC, Quantitative phosphoproteomic analysis of plasma membrane proteins reveals regulatory mechanisms of plant innate immune responses. Plant J, 2007. 51(5): p. 931–40.

6. Withers, J and Dong, X, Post-translational regulation of plant immunity. Curr Opin Plant Biol, 2017. 38: p. 124–132.

7. Tena, G, Boudsocq, M, and Sheen, J, Protein kinase signaling networks in plant innate immunity. Curr Opin Plant Biol, 2011. 14(5): p. 519–29.

8. Lu, D, Lin, W, Gao, X, Wu, S, Cheng, C, Avila, J, Heese, A, Devarenne, TP, He, P, and Shan, L, Direct Ubiquitination of Pattern Recognition Receptor FLS2 Attenuates Plant Innate Immunity. 2011. 332(6036): p. 1439–1442.

9. Feng, B, Liu, C, de Oliveira, MVV, Intorne, AC, Li, B, Babilonia, K, de Souza Filho, GA, Shan, L, and He, P, Protein Poly(ADP-ribosyl)ation Regulates Arabidopsis Immune Gene Expression and Defense Responses. PLOS Genetics, 2015. 11(1): p. e1004936.

10. Macho, AP and Zipfel, C, Plant PRRs and the activation of innate immune signaling. Mol Cell, 2014. 54(2): p. 263–72.

11. Yu, X, Feng, B, He, P, and Shan, L, From Chaos to Harmony: Responses and Signaling upon Microbial Pattern Recognition. Annu Rev Phytopathol, 2017. 55: p. 109–137.

12. Huffaker, A, Pearce, G, and Ryan, CA, An endogenous peptide signal in Arabidopsis activates components of the innate immune response. Proc Natl Acad Sci U S A, 2006. 103(26): p. 10098–103.

13. Huffaker, A, Dafoe, NJ, and Schmelz, EA, ZmPep1, an ortholog of Arabidopsis elicitor peptide 1, regulates maize innate immunity and enhances disease resistance. Plant Physiol, 2011. 155(3): p. 1325–38.

14. Huffaker, A, Pearce, G, Veyrat, N, Erb, M, Turlings, TC, Sartor, R, Shen, Z, Briggs, SP, Vaughan, MM, Alborn, HT, Teal, PE, and Schmelz, EA, Plant elicitor peptides are conserved signals regulating direct and indirect antiherbivore defense. Proc Natl Acad Sci U S A, 2013. 110(14): p. 5707–12.

15. Trivilin, AP, Hartke, S, and Moraes, MG, Components of different signalling pathways regulated by a new orthologue of AtPROPEP1 in tomato following infection by pathogens. 2014. 63(5): p. 1110–1118.

16. Lee, MW, Huffaker, A, Crippen, D, Robbins, RT, and Goggin, FL, Plant elicitor peptides promote plant defences against nematodes in soybean. Mol Plant Pathol, 2018. 19(4): p. 858–869.

17. Ruiz, C, Nadal, A, Montesinos, E, and Pla, M, Novel Rosaceae plant elicitor peptides as sustainable tools to control Xanthomonas arboricola pv. pruni in Prunus spp. Mol Plant Pathol, 2018. 19(2): p. 418–431.

18. Lori, M, van Verk, MC, Hander, T, Schatowitz, H, Klauser, D, Flury, P, Gehring, CA, Boller, T, and Bartels, S, Evolutionary divergence of the plant elicitor peptides (Peps) and their receptors: interfamily incompatibility of perception but compatibility of downstream signalling. J Exp Bot, 2015. 66(17): p. 5315–25.

19. Hander, T, Fernandez-Fernandez, AD, Kumpf, RP, Willems, P, Schatowitz, H, Rombaut, D, Staes, A, Nolf, J, Pottie, R, Yao, P, Goncalves, A, Pavie, B, Boller, T, Gevaert, K, Van Breusegem, F, Bartels, S, and Stael, S, Damage on plants activates Ca(2+)-dependent metacaspases for release of immunomodulatory peptides. Science, 2019. 363(6433).

20. Yamaguchi, Y, Pearce, G, and Ryan, CA, The cell surface leucine-rich repeat receptor for AtPep1, an endogenous peptide elicitor in Arabidopsis, is functional in transgenic tobacco cells. Proc Natl Acad Sci U S A, 2006. 103(26): p. 10104–9.

21. Yamaguchi, Y, Huffaker, A, Bryan, AC, Tax, FE, and Ryan, CA, PEPR2 is a second receptor for the Pep1 and Pep2 peptides and contributes to defense responses in Arabidopsis. Plant Cell, 2010. 22(2): p. 508–22.

22. Krol, E, Mentzel, T, Chinchilla, D, Boller, T, Felix, G, Kemmerling, B, Postel, S, Arents, M, Jeworutzki, E, Al-Rasheid, KA, Becker, D, and Hedrich, R, Perception of the Arabidopsis danger signal peptide 1 involves the pattern recognition receptor AtPEPR1 and its close homologue AtPEPR2. J Biol Chem, 2010. 285(18): p. 13471–9.

23. Tintor, N, Ross, A, Kanehara, K, Yamada, K, Fan, L, Kemmerling, B, Nurnberger, T, Tsuda, K, and Saijo, Y, Layered pattern receptor signaling via ethylene and endogenous elicitor peptides during Arabidopsis immunity to bacterial infection. Proc Natl Acad Sci U S A, 2013. 110(15): p. 6211–6.

24. Ross, A, Yamada, K, Hiruma, K, Yamashita-Yamada, M, Lu, X, Takano, Y, Tsuda, K, and Saijo, Y, The Arabidopsis PEPR pathway couples local and systemic plant immunity. Embo j, 2014. 33(1): p. 62–75.

25. Postel, S, Kufner, I, Beuter, C, Mazzotta, S, Schwedt, A, Borlotti, A, Halter, T, Kemmerling, B, and Nurnberger, T, The multifunctional leucine-rich repeat receptor kinase BAK1 is implicated in Arabidopsis development and immunity. Eur J Cell Biol, 2010. 89(2-3): p. 169–74.

26. Liu, Z, Wu, Y, Yang, F, Zhang, Y, Chen, S, Xie, Q, Tian, X, and Zhou, JM, BIK1 interacts with PEPRs to mediate ethylene-induced immunity. Proc Natl Acad Sci U S A, 2013. 110(15): p. 6205–10.

27. Lu, D, Wu, S, Gao, X, Zhang, Y, Shan, L, and He, P, A receptor-like cytoplasmic kinase, BIK1, associates with a flagellin receptor complex to initiate plant innate immunity. Proc Natl Acad Sci U S A, 2010. 107(1): p. 496–501.

28. Li, L, Li, M, Yu, L, Zhou, Z, Liang, X, Liu, Z, Cai, G, Gao, L, Zhang, X, Wang, Y, Chen, S, and Zhou, JM, The FLS2-associated kinase BIK1 directly phosphorylates the NADPH oxidase RbohD to control plant immunity. Cell Host Microbe, 2014. 15(3): p. 329–38.

29. Lal, NK, Nagalakshmi, U, Hurlburt, NK, Flores, R, Bak, A, Sone, P, Ma, X, Song, G, Walley, J, Shan, L, He, P, Casteel, C, Fisher, AJ, and Dinesh-Kumar, SP, The Receptor-like Cytoplasmic Kinase BIK1 Localizes to the Nucleus and Regulates Defense Hormone Expression during Plant Innate Immunity. Cell Host Microbe, 2018. 23(4): p. 485–497.e5.

30. Monaghan, J, Matschi, S, Shorinola, O, Rovenich, H, Matei, A, Segonzac, C, Malinovsky, FG, Rathjen, JP, MacLean, D, Romeis, T, and Zipfel, C, The calcium-dependent protein kinase CPK28 buffers plant immunity and regulates BIK1 turnover. Cell Host Microbe, 2014. 16(5): p. 605–15.

31. Wang, J, Grubb, LE, Wang, J, Liang, X, Li, L, Gao, C, Ma, M, Feng, F, Li, M, Li, L, Zhang, X, Yu, F, Xie, Q, Chen, S, Zipfel, C, Monaghan, J, and Zhou, JM, A Regulatory Module Controlling Homeostasis of a Plant Immune Kinase. Mol Cell, 2018. 69(3): p. 493–504.e6.

32. Walley, JW, Sartor, RC, Shen, Z, Schmitz, RJ, Wu, KJ, Urich, MA, Nery, JR, Smith, LG, Schnable, JC, Ecker, JR, and Briggs, SP, Integration of omic networks in a developmental atlas of maize. Science, 2016. 353(6301): p. 814–8.

33. Golovkin, M and Reddy, AS, An SC35-like protein and a novel serine/arginine-rich protein interact with Arabidopsis U1-70K protein. J Biol Chem, 1999. 274(51): p. 36428–38.

34. Carvalho, RF, Szakonyi, D, Simpson, CG, Barbosa, IC, Brown, JW, Baena-Gonzalez, E, and Duque, P, The Arabidopsis SR45 Splicing Factor, a Negative Regulator of Sugar Signaling, Modulates SNF1-Related Protein Kinase 1 Stability. Plant Cell, 2016. 28(8): p. 1910–25.

35. Zhang, XN, Shi, Y, Powers, JJ, Gowda, NB, Zhang, C, Ibrahim, HMM, Ball, HB, Chen, SL, Lu, H, and Mount, SM, Transcriptome analyses reveal SR45 to be a neutral splicing regulator and a suppressor of innate immunity in Arabidopsis thaliana. BMC Genomics, 2017. 18(1): p. 772.

36. Asai, T, Tena, G, Plotnikova, J, Willmann, MR, Chiu, WL, Gomez-Gomez, L, Boller, T, Ausubel, FM, and Sheen, J, MAP kinase signalling cascade in Arabidopsis innate immunity. Nature, 2002. 415(6875): p. 977–83.

37. Zhang, M, Su, J, Zhang, Y, Xu, J, and Zhang, S, Conveying endogenous and exogenous signals: MAPK cascades in plant growth and defense. Curr Opin Plant Biol, 2018. 45(Pt A): p. 1–10.

38. Mei, Y, Zhang, C, Kernodle, BM, Hill, JH, and Whitham, SA, A Foxtail mosaic virus Vector for Virus-Induced Gene Silencing in Maize. Plant Physiol, 2016. 171(2): p. 760–72.

39. Anko, ML, Regulation of gene expression programmes by serine-arginine rich splicing factors. Semin Cell Dev Biol, 2014. 32: p. 11–21.

40. Jeong, S, SR Proteins: Binders, Regulators, and Connectors of RNA. Mol Cells, 2017. 40(1): p. 1–9.

41. Thines, B, Katsir, L, Melotto, M, Niu, Y, Mandaokar, A, Liu, G, Nomura, K, He, SY, Howe, GA, and Browse, J, JAZ repressor proteins are targets of the SCF(COI1) complex during jasmonate signalling. Nature, 2007. 448(7154): p. 661–5.

42. Chung, HS and Howe, GA, A critical role for the TIFY motif in repression of jasmonate signaling by a stabilized splice variant of the JASMONATE ZIM-domain protein JAZ10 in Arabidopsis. Plant Cell, 2009. 21(1): p. 131–45.

43. Jabs, T, Tschöpe, M, Colling, C, Hahlbrock, K, and Scheel, DJPotNAoS, Elicitor-stimulated ion fluxes and O2– from the oxidative burst are essential components in triggering defense gene activation and phytoalexin synthesis in parsley. 1997. 94(9): p. 4800–4805.

44. Acharya, BR, Raina, S, Maqbool, SB, Jagadeeswaran, G, Mosher, SL, Appel, HM, Schultz, JC, Klessig, DF, and Raina, R, Over expression of CRK13, an Arabidopsis cysteine-rich receptor-like kinase, results in enhanced resistance to Pseudomonas syringae. Plant J, 2007. 50(3): p. 488–99.

45. Waese, J, Fan, J, Pasha, A, Yu, H, Fucile, G, Shi, R, Cumming, M, Kelley, LA, Sternberg, MJ, Krishnakumar, V, Ferlanti, E, Miller, J, Town, C, Stuerzlinger, W, and Provart, NJ, ePlant: Visualizing and Exploring Multiple Levels of Data for Hypothesis Generation in Plant Biology. Plant Cell, 2017. 29(8): p. 1806–1821.

46. Ali, GS, Palusa, SG, Golovkin, M, Prasad, J, Manley, JL, and Reddy, AS, Regulation of plant developmental processes by a novel splicing factor. PLoS One, 2007. 2(5): p. e471.

47. Liese, A and Romeis, T, Biochemical regulation of in vivo function of plant calcium-dependent protein kinases (CDPK). Biochim Biophys Acta, 2013. 1833(7): p. 1582–9.

48. Klimecka, M and Muszynska, G, Structure and functions of plant calcium-dependent protein kinases. Acta Biochim Pol, 2007. 54(2): p. 219–33.

49. Manley, JL and Tacke, R, SR proteins and splicing control. Genes Dev, 1996. 10(13): p. 1569–79.

50. Xing, D, Wang, Y, Hamilton, M, Ben-Hur, A, and Reddy, AS, Transcriptome-Wide Identification of RNA Targets of Arabidopsis SERINE/ARGININE-RICH45 Uncovers the Unexpected Roles of This RNA Binding Protein in RNA Processing. Plant Cell, 2015. 27(12): p. 3294–308.

